# Genomic epidemiology of *Treponema pallidum* and circulation of strains with diminished *tprK* antigen variation capability in Seattle, 2021-2022

**DOI:** 10.1101/2023.05.12.540601

**Authors:** Nicole A.P. Lieberman, Carlos Avendaño, Shah A. K. Mohamed Bakhash, Ethan Nunley, Hong Xie, Lorenzo Giacani, Anna Berzkalns, Olusegun O. Soge, Tara B. Reid, Matthew R. Golden, Alexander L. Greninger

**Author notes:** **Corresponding author contact:** Alexander L Greninger, 850 Republican St Rm S130, Seattle WA 98118.

## Abstract

**Background:** Syphilis incidence continues to increase dramatically in the United States and yet little is known about *Treponema pallidum* (TP) genomic epidemiology within American metropolitan areas.

**Methods:** We performed whole genome sequencing and *tprK* deep sequencing of 28 TP-containing specimens collected mostly from remnant Aptima swabs from 24 individuals from Seattle Sexual Health Clinic during 2021-2022.

**Results:** All 12 individuals infected with Nichols lineage strains were MSM, while a specific SS14 cluster (average 0.33 SNPs) included 1 MSW and five women. All TP strains sequenced were azithromycin resistant via 23S rRNA A2058G mutation. Identical TP genomic sequences were found in pharyngeal and rectal swab specimens taken from the same individuals concurrently. *tprK* sequences were less variable between patient-matched specimens and between epidemiologically-linked clusters. We detected a 528 bp deletion in the *tprK* donor site locus, eliminating nine *tprK* donor sites, in TP genomes of three individuals with secondary syphilis, associated with diminution of overall *tprK* sequence diversity.

**Conclusions:** We developed an end-to-end workflow for public health genomic surveillance of TP from remnant Aptima swab specimens. With its high rate of gene conversion, *tprK* sequencing may assist in linking cases beyond routine TP genome sequencing. TP strains with deletions in *tprK* donor sites currently circulate and are associated with diminished antigenic diversity of the TprK putative outer membrane protein.

## Introduction

The incidence of syphilis in the United States has increased for the last two decades and remains a major public health concern [1]. Among all individuals, the incidence of syphilis in the United States was 16.2 cases per 100,000 persons in 2021, with higher incidence among vulnerable populations including men who have sex with men (MSM) and persons experiencing homelessness [1,2]. Notably, despite routine screening during prenatal care, the incidence of congenital syphilis has risen more than 219% from 2017-2021 to 77.9 cases per 100,000 live births [1]. Improved diagnostics, public health surveillance including genomic epidemiology, and outreach to and education of vulnerable and general populations are all necessary tools to combat this epidemic.

Syphilis is caused by the spirochete bacterium *Treponema pallidum* subspecies *pallidum* (TP). Few putative outer membrane proteins are encoded within its small (1.13 Mb) genome, and of those, most appear to be poorly and not consistently expressed on the bacterial surface throughout its life cycle [3–5], facilitating evasion of the adaptive immune system. An additional, and likely more important, immune evasion strategy employed by TP involves continuous antigenic variation of seven predicted surface-exposed variable (V) loop regions of the putative TprK outer membrane porin by non-reciprocal recombination of DNA sequences from donor sites (DS) flanking the *tprD* (*tp0131*) locus [6–10]. Recent studies in rabbits using a TP strain genetically modified to ablate 96% of donor sites demonstrated that the inability to vary the antigen composition of TprK leads to reduced V region diversity, attenuated disease manifestations, and decreased ability of TP to persist in the host, highlighting the importance of this virulence factor in pathogen persistence [11].

Despite an increase in global TP sequencing efforts over the past 5 years, significantly less genomic data is available for this bacterial pathogen compared to many others due to the paucity of molecular testing and, to date, none of these sequencing efforts has described the genomic epidemiology focused on an American metropolitan area [12–16]. Here, using a novel workflow for public health genomics from mostly remnant Aptima swabs, we performed whole genome sequencing (WGS) and *tprK* deep sequencing of 28 swab specimens from 24 individuals attending the Seattle Sexual Health Clinic during 2021-2022.

## Materials and Methods

### Patient specimens and TP detection

Specimens came from two studies conducted in the Seattle King County Sexual Health Clinic. Both studies were approved by the University of Washington IRB and participants submitted written informed consent. The first involves testing remnant rectal, pharyngeal, urine, vaginal, and serum specimens obtained from patients seeking clinic care in the clinic using an Aptima-based experimental transcription mediated assay (TMA). The second study included swabs from anogenital lesions (if present) and oropharyngeal mucosa from patients with active syphilis collected in 10mM Tris-HCl, 0.1M EDTA, and 0.5% SDS lysis buffer. Samples were collected from approximately February 2021 to May 2022.

DNA was purified from 200μL Aptima tube specimens using the MagNA Pure Total Nucleic Acid Isolation Kit I (Roche Applied Science, Indianapolis, IN) external lysis protocol on the automated MagNA Pure 24 Instrument and 50µL elution. Swabs from non-TMA positive specimens were collected in 1mL of 10mM Tris-HCl, 0.1 M EDTA, 0.5% SDS and extraction was performed on 200µL sample buffer using QIAamp DNA mini kit (Qiagen, Valencia, CA) with a 100µL elution.

### Tp47 quantification

Treponemal load quantification was performed using TaqMan qPCR targeting the *tp47* locus and multiplexed with human beta-globin as previously described [15]. 2μL DNA was amplified in a 30μL reaction consisting of 14.33μL QuantiTect Multiplex PCR NoROX mix (Qiagen), 0.67μL QuantiTect Multiplex PCR mix with ROX (Qiagen), 0.03μL UNG (Lucigen), 0.1μL EXO for internal control, 0.075μL each 100μM forward (5’-CAAGTACGAGGGGAACATCGAT-3’) and reverse (5’-TGATCGCTGACAAGCTTAGG-3’) primers, and 0.03μL 100μM probe (6FAM-CGGAGACTCTGATGGATGCTGCAGTT-NFQMGB). qPCR was performed on a QuantStudio7 with 50°C 2min, 95°C 15min, followed by 45 cycles of 94°C 1min and 60°C 1min [17]. A gblock standard carrying the target amplicon was used to determine the copy number per microliter.

### TP genome library generation and sequencing

TP pre-capture libraries were prepared as previously described [15] using the KAPA HyperPlus kit with UDI adapters (Roche). 17.5µL DNA was used as input for fragmentation for eight minutes, amplified with 14 cycles of PCR, and cleaned with 0.8X AMPureXP beads (Beckman Coulter). Three libraries totaling 250-400ng were pooled together based on treponemal load and were enriched using custom biotinylated probes based on NC_010741.1 and the IDT xGen Hybridization and Wash kit with a 4-hour hybridization followed by 14 cycles of post-capture PCR. Enriched libraries were purified using a 0.8X AMPureXP cleanup. Sequencing was performed on 2x151bp Illumina NextSeq2000 or NovaSeq6000 runs.

### Genome assembly

Fastqs were processed and genomes assembled using a custom pipeline, available at https://github.com/greninger-lab/T.Pallidum-Seattle. Briefly, paired-end reads were adapter- and quality-trimmed by Trimmomatic v0.35 [18], kmer-filtered to match TP by bbduk v38.86 [19], then mapped to the TP SS14 reference genome (NC_021508.1) using Bowtie2 v2.4.1 [20] with default parameters followed by deduplication using Picard v2.23.3. Separately, de novo assembly was performed with Unicycler v0.4.4 [21] using reads filtered with bbduk [19] to remove repetitive regions of the genome, including the repeat regions of the *arp* and *TP0470* genes, as well as *tprC/D/F/I* and *tprE/G/J* loci. A hybrid fasta combining reference mapping and de novo sequences was generated and deduplicated reads were remapped to this hybrid using default Bowtie2 settings, local misalignments corrected with Pilon v1.23.0 [22], and a final Bowtie2 remapping to the Pilon consensus used as input to a custom R script to close gaps and generate a final consensus sequence. Raw data files and consensus genomes have been deposited to SRA and GenBank under BioProject PRJNA961304 (Table S1 in Supplemental Data).

### Phylogenetic analysis

Genomes masked at repetitive loci [15] were aligned with MAFFT v7.490 and iqtree v2.0.3 used to generate a whole genome maximum likelihood phylogeny using 1000 ultrafast bootstraps and automated selection of the best substitution model.

### TprK sequencing

Specimens were run in technical duplicates for *tprK* deep sequencing using 10.5µL DNA added to 12.5µL 2X CloneAmp Pre-mix (Takara), 1µL 10µM barcoded tprK-F and 1µL 10µM barcoded tprK-R (Table S2 in Supplemental Data). PCR conditions were 98°C 2 min; 35x cycles 98°C 10s, 62°C 15s, 72°C 90s; 72°C 5 min; 4°C hold. 12.5µL of PCR product was cleaned using 0.6X AmpureXP and eluted in 20µL water. 15µL cleaned PCR product was used as input for the Illumina DNA Prep(M) library preparation with 12 cycles of dual-index PCR. 12.5µL amplified library was cleaned using 0.75X AmpureXP, eluted in 20µL water. Sequencing was performed on 2x151 bp Illumina NextSeq2000 or NovaSeq6000 runs.

### Donor Site Deletion PCR and Sanger Sequencing

TP deletion PCR with input ranging between 500 - 1,000 total copies in a 25 μL reaction volume including 10.5µL template, 12.5µL 2X CloneAmp Pre-Mix (Takara) and 1µL each 10µM of tpdel-F (5’-TGCTCTCGCATGGCTATGTT-3’) and tpdel-R (5’-GCGCGGAAATAAGACGAACG-3’). Conditions were 98°C 2 min; 35x cycles 98°C 10s, 57°C 15s, 72°C 30s; 72°C 5 min; 10°C hold. 2µL PCR product was run on a 1.2% agarose gel to confirm band sizes. PCR products were cleaned using 1.8X AmpureXP and sequencing confirmed via Sanger sequencing using the same primers as for PCR.

### tprK bioinformatic analysis

Paired-end *tprK* reads were adapter-trimmed and merged in bbmerge [19] using maximum stringency and processed with our custom pipeline [16] to identify V regions (https://github.com/greninger-lab/T.Pallidum-Seattle). Sequences that resulted in frameshift or nonsense mutations were discarded. In-frame reads were randomly downsampled to an equivalent number of sequences (3471) per V region per replicate. Concordance between replicates was confirmed by determining the Pearson correlation coefficient between replicates exceeded 0.85, then reads per V region averaged across both replicates. Sequences only appearing only in a single replicate or below the sequencing error threshold (0.5%, 17 reads) were discarded. High-confidence short reads were then locally aligned using BLAST v2.13.0 [23] to the sample-matched donor site locus determined during WGS, using a word size 10 and 100% identity. The top BLAST hit for each V region was then annotated as belonging to one of 53 known donor sites [9].

### Statistics and visualization

All analyses were performed in R v4.2.3 using packages ggplot2 v3.4.2 [24] and ggtree v3.7.2 [25].

## Results

### Descriptive epidemiology of sequenced T. pallidum specimens

As part of a clinical study of TP testing in King County, we had available 86 oropharyngeal and rectal swabs in Aptima buffer positive for TP by Hologic real-time transcription mediated amplification [26,27]. Of the 86 positive Aptima swabs, 14 specimens did not test positive by *tp47* qPCR due to low treponemal load. Among samples positive by both *tp47* qPCR and TMA, comparison of the qPCR-derived treponemal loads with rTMA time revealed a robust negative correlation (r = −0.78), with successful genome recovery for specimens for which >600 genome copies were available for capture library preparation using a 17.5µL input volume (Figure 1, Table S3).

**Figure 1.**
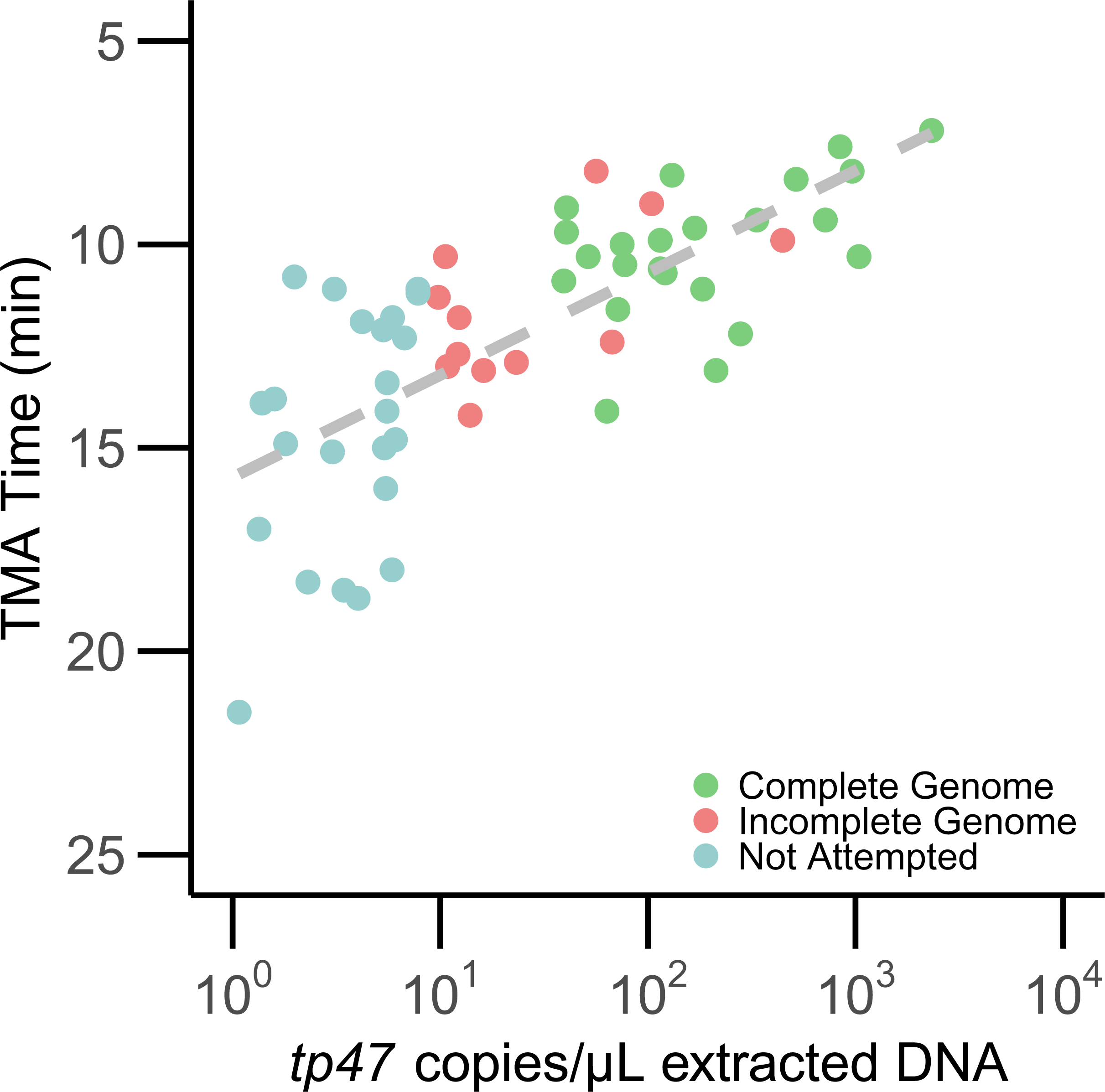
Relationship between TP genome copies in extracted DNA, rTMA time on Aptima assay, and ability to recover complete genome. TP genome copies as determined by *tp47* qPCR were highly correlated with rTMA time taken directly from the Hologic Aptima assay (r=-0.78). Complete genomes were defined as having <5.3% missing data in the genome (green), while incomplete genomes had >5.3% missing data (red). We did not attempt capture on specimens that had <10 copies/µL extracted DNA based on prior experience (light blue).

We initially attempted hybridization capture sequencing on 41 specimens with treponemal loads >10 *tp47* copies/µL extracted genomic DNA, consisting of DNA from 35 Aptima and 6 non-Aptima swabs. Twenty-eight near complete TP genomes (<5.3% missing data) were recovered from 24 individuals (Table 1). Patients were 75% male, with 89% of males MSM. Age ranged between 20-64 years old, with the most common age range 30-34 years old (25% of patients). Eighty percent of patients had a clinical diagnosis of primary or secondary syphilis. None were HIV positive.

**Table1.** Demographic and Clinical Summary for 24 Individuals for which a TP Genome was Recovered

| **Sex** | **N (%)** |
| --- | --- |
| Male | 18 (75%) |
| Female | 6 (25%) |
| **MSM (males only)** |  |
| ☐☐☐☐ Yes | 16 (89%) |
| ☐☐☐☐ No | 2 (11%) |
| **Age Category (years)** |  |
| 20-24 | 3 (12%) |
| 25-29 | 4 (17%) |
| 30-34 | 6 (25%) |
| 35-44 | 5 (21%) |
| 45-54 | 4 (17%) |
| 55-64 | 2 (8%) |
| **Race/Ethnicity** |  |
| White | 12 (50%) |
| Asian | 3 (12%) |
| Black | 2 (8%) |
| Hispanic | 5 (21%) |
| NHPI | 1 (4%) |
| Unknown | 1 (4%) |
| **Disease Stage** |  |
| Primary | 10 (42%) |
| Secondary | 9 (38%) |
| Early latent | 2 (8%) |
| Late or Unknown duration | 3 (12%) |
| **IV Drug Use in Past Year** |  |
| Yes | 6 (25%) |
| No | 3 (12%) |
| Unknown | 15 (62%) |
| **Meth Use in Past Year** |  |
| Yes | 7 (29%) |
| No | 11 (46%) |
| Unknown | 6 (25%) |
| **Sample Swab Location (total N=28 swabs)** |  |
| Throat | 13 (46%) |
| Rectal | 7 (25%) |
| Other Lesion | 4 (14%) |
| Vaginal | 4 (14%) |

### Phylogenetic clustering of TP genomes associated with likely transmission chains

Phylogenetic analysis revealed that 15 genomes (representing 12 individuals) belonged to the Nichols lineage, while the remaining 13 genomes (12 individuals) were from the SS14 lineage (Figure 2; Table S3 in Supplemental Data). All strains were azithromycin resistant via the A2058G mutation in the 23S rRNA locus. All four patient-matched sample pairs had identical genomes outside of *tprK*. Disease stage was significantly associated with lineage (Chi-square, *p*=0.0027). Nichols lineage strains were more likely to be obtained from patients with secondary syphilis (n= 9, 89% Nichols vs 11% SS14), and SS14 lineage strains more likely to be obtained from patients with primary syphilis (n=10, 20% Nichols vs 80% SS14).

**Figure 2.**
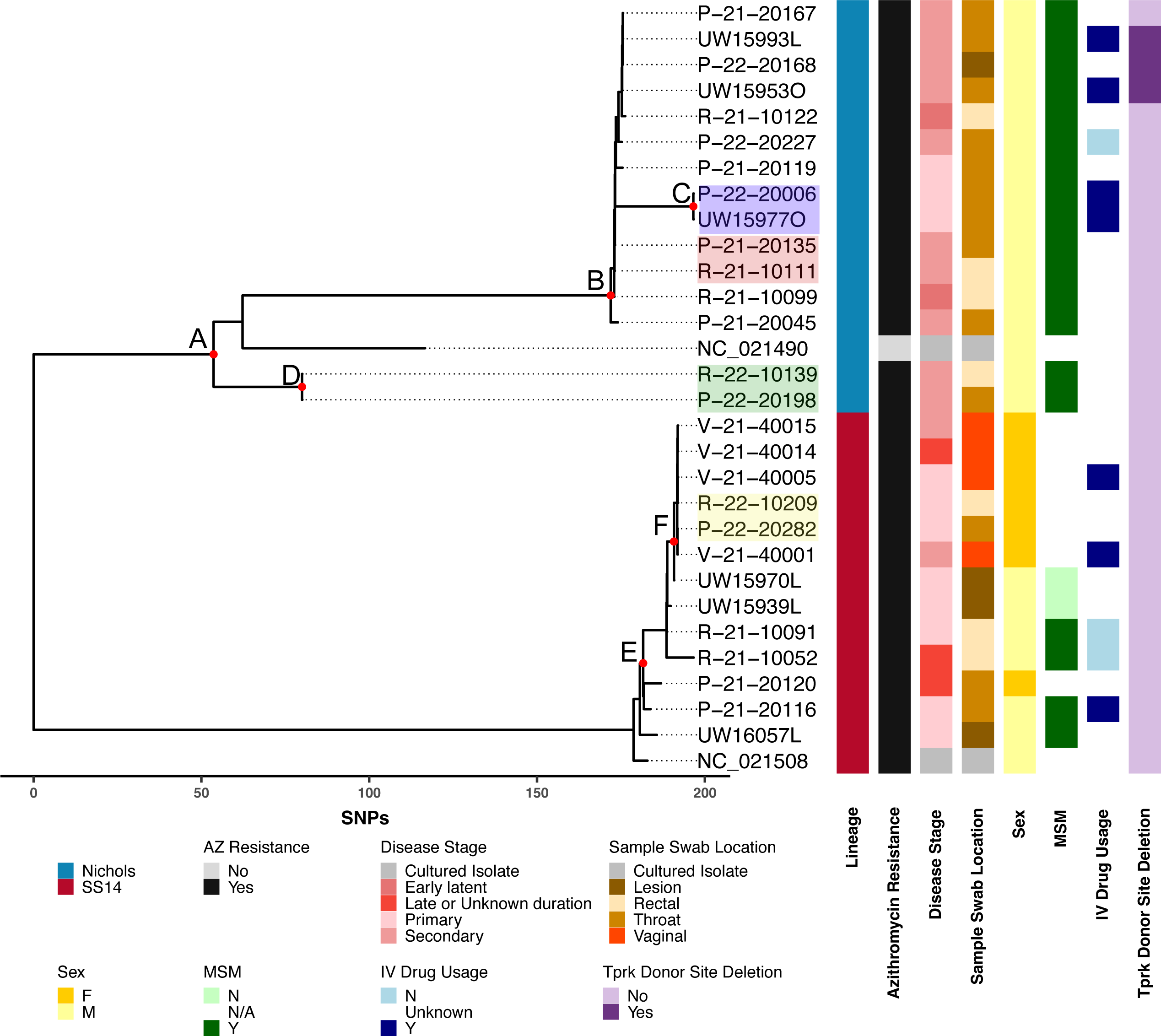
Maximum likelihood phylogeny of TP strains in Seattle from 2021-2022. Red nodes labeled with letters indicate high-confidence bootstrap values > 0.9. Strains sequenced from paired pharyngeal (P,O) and rectal (R) swabs taken at the same time from three separate individuals are back highlighted in pink, green, and yellow. Node C (P−22−20006 and UW15977O) consists of paired pharyngeal swab samples from the same individual on the same day and is highlighted in purple. TP reference sequences for Nichols (NC_021490) and SS14 (NC_021508) are used for orientation.

Notably, we found that all 12 individuals with Nichols-lineage samples were MSM, consistent with data from Japan and our prior global sequencing efforts [15,28], though contrasting with recent genetic surveillance in Amsterdam [29]. In contrast, only four of the twelve individuals with SS14 samples were MSM; the remainder were MSW (two) and females (six) (Chi-square, *p*=0.00053).

Within the SS14 lineage, a cluster comprising five of the six female patients and one MSW (node F) separated by a median of 0 single nucleotide variants (SNVs, average 0.33 SNVs) is suggestive of local circulation of a single SS14-lineage strain among heterosexual individuals. Strains comprising node F were separated from the SS14 strains derived from MSM (R-21-10091, R-21-20052, P-21-20116, and UW16057L) by an average of 10.2 SNVs (median= 11, range 2-16 SNVs) and from the other female-derived strain (P-21-20120) by 14 or 15 SNVs, which itself was separated from the four MSM strains by a median 11.5 SNVs (range 7-20 SNVs). The four MSM strains differed each other by a median 11 SNVs (range 8-21 SNVs). However, low diversity and/or undersampling limits the interpretability of phylogenetic and epidemiologic relationships between the node F cluster and the other SS14 strains derived from MSM individuals.

Within the Nichols lineage, very limited diversity was seen, with a median number of 2 SNVs (average 6.3 SNVs, Table S4 in Supplemental Data) separating strains within node B (range 0-26 SNVs). Interestingly, strains comprising node D (R-22-10139 and R-22-20198) taken from the same individual were more distantly related to other Nichols genomes sequenced here (average of 140.5 SNVs), and were most closely related to Nichols subclades C and D [as defined in 15], though would likely constitute a separate Nichols subclade based on genetic distance.

### Correlated tprK sequences associated with likely transmission chains

We next performed deep sequencing of the *tprK* gene to examine V region composition and donor site usage by strain. We were particularly interested in how similar the pattern of use of V region sequences were within epidemiologically-linked local clusters and in patients with samples taken from different sites at the same time. We first examined potential confounders and found no relationship between the number of unique high-confidence V regions per sample and the number of TP genome copies input to the *tprK* library prep replicates (Supplemental Fig 1A, Pearson coefficient r= 0.204, *p*=0.270). There was also no difference in the number of unique V regions by lineage or clinical stage (Supplemental Figure 2B, *p*>0.05, Welch’s t-test), despite previous studies of full-length haplotypes showing increased diversity of *tprK* in secondary samples [9,30].

We hypothesized that epidemiologically-linked TP strains would have a more similar V region composition than distantly related strains. Because traditional phylogenetic methods reconstruct relationships based on single SNP evolution, they are not suitable to compare sequences from *tprK*, which changes its seven variable regions via recombination of up to tens of bases at a time. Instead, we calculated the correlation between the proportions of *tprK* V region sequences between sample pairs, epidemiologically-linked node F, and unlinked samples in the same or opposite lineage. As expected, we found high (median 0.93) correlation coefficients between paired samples from the same individual (Figure 3A-B), including a matched sample (UW15993O) from a *tprK* donor site deletion strain (UW15993L) for which a whole genome could not be recovered. Although lower than for paired samples, we found that the likely heterosexual cluster defined by node F (Figure 2) had significantly higher pairwise Pearson coefficients than between more distantly related samples either in the same or the opposite lineage (*p*<0.0001, One-way ANOVA with Tukey’s HSD). This is strongly suggestive of close relationships among samples in the cluster, though the background rate of *tprK* gene conversion in human syphilis infection is unknown.

**Figure 3:**
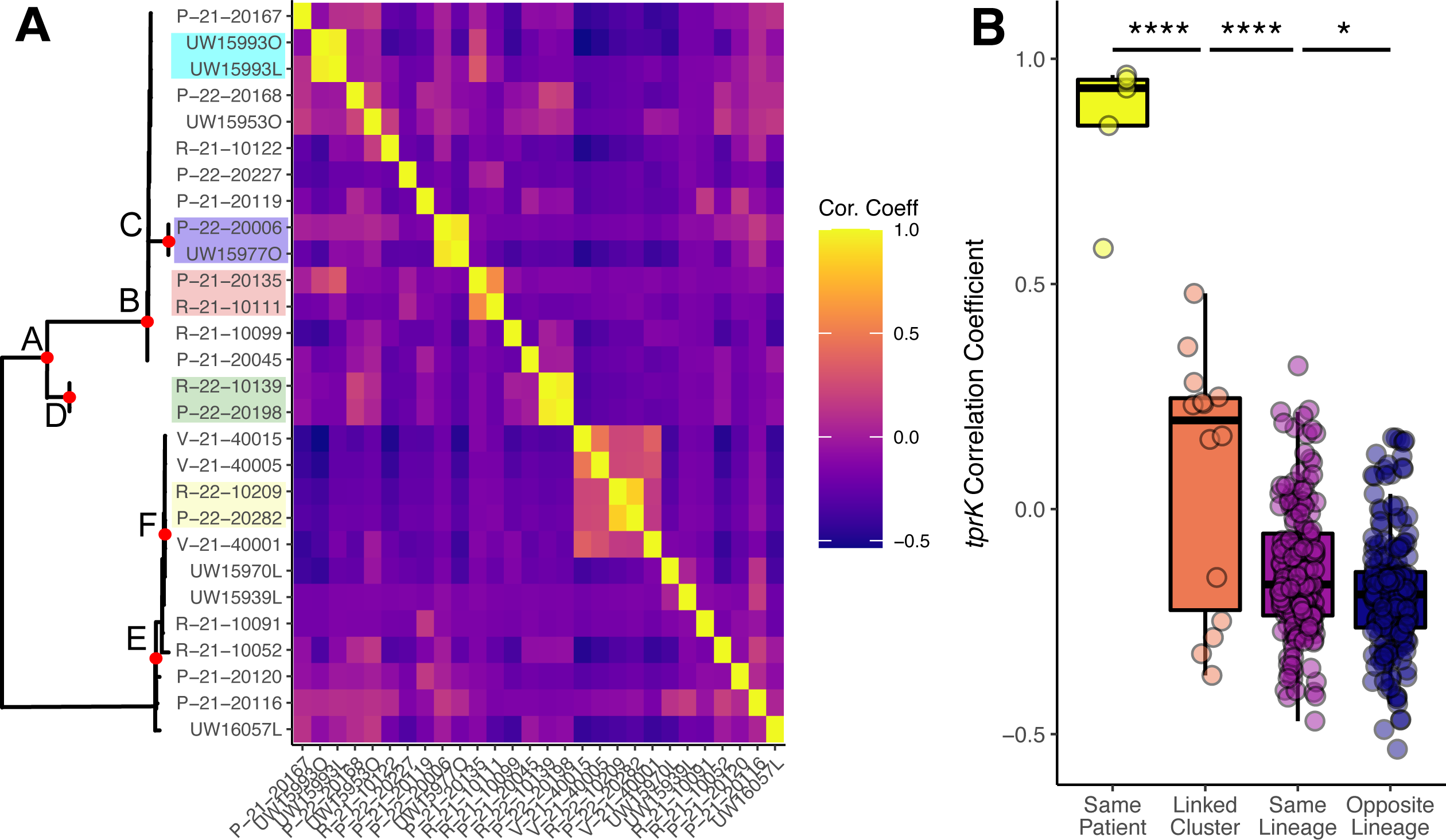
*tprK* V region sequence proportion correlations from strains in this study. A) *tprK* V region sequences and proportions were compared pairwise and Pearson’s correlation coefficient determined. More similar patterns of V region use result in a higher coefficient. The outline of the phylogeny from Figure 2 (slightly modified to only include strains yielding *tprK* sequence data) is shown for orientation. Patient-matched samples are shown in the same color. B) Pearson coefficients are significantly different among patient-matched, Node F cluster-associated, and within the same or opposite lineages (\*\*\*\**p*<0.0001, \**p*<0.05, One-way ANOVA with Tukey’s HSD).

### Identification of a large deletion in tprK donor site locus associated with reduced tprK diversity

Intriguingly, three Nichols specimens (P-21-20167, P-22-20168 and UW15993L; Figure 2) contained a 528bp deletion, bounded by a 16 bp direct repeat, within the donor site locus for *tprK*, resulting in loss of nine of the 53 known *tprK* donor sites (Figure 4). The deletion was confirmed by PCR band size (Supplemental Figure 2) and Sanger sequencing. There were no differences in the TP genome sequence between the three samples, strongly suggestive of transmission of a single deletion-bearing strain rather than independent deletion events. Although experimental elimination of 96% of donor sites results in reduced *tprK* diversity, severe attenuation of early lesions, and diminished ability to persist in the rabbit host [11], ablation of only nine donor sites did not appear to prevent lesion appearance or prevent transmission in humans as these three patients had clinically evident secondary syphilis.

**Figure 4:**
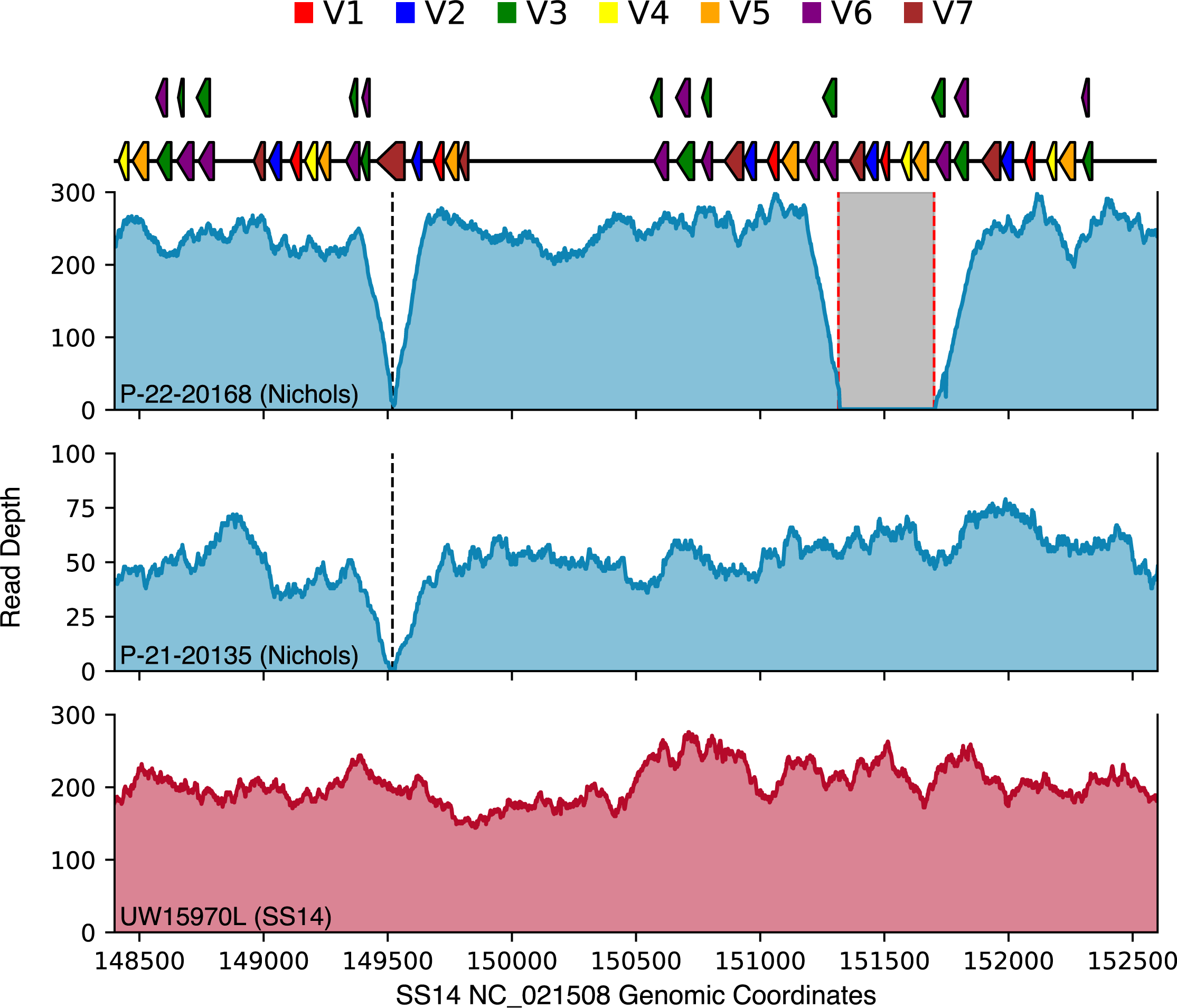
Nichols lineage strain with a 528 bp deletion in *tprK* donor site locus. Sequencing read depth is depicted across the *tprK* donor site locus for three TP strains sequenced in this study, using SS14 genome reference NC_021508.1. The 528 bp deletion that removes nine donor sites is denoted by red dotted lines in strain P-22-20168. A 51 bp deletion present in all sequenced Nichols strains is depicted in strain P-21-20135. SS14-lineage strains have neither deletion, as indicated by sequencing depth for strain UW15970L.

We next examined V region composition between strains with and without the *tprK* donor site deletion. Due to the potential confounding effects of lineage and stage on the analysis, we compared the total V region diversity (number of unique V regions recovered) from deletion strains only to the diversity in other Nichols secondary syphilis samples. Strains harboring the *tprK* donor site deletion had significantly reduced diversity (Figure 5A, *p*<0.05, Welch’s t-test), consistent with the phenotype seen in rabbits infected with TP with t*prK* donor sites knocked out [11].

**Figure 5:**
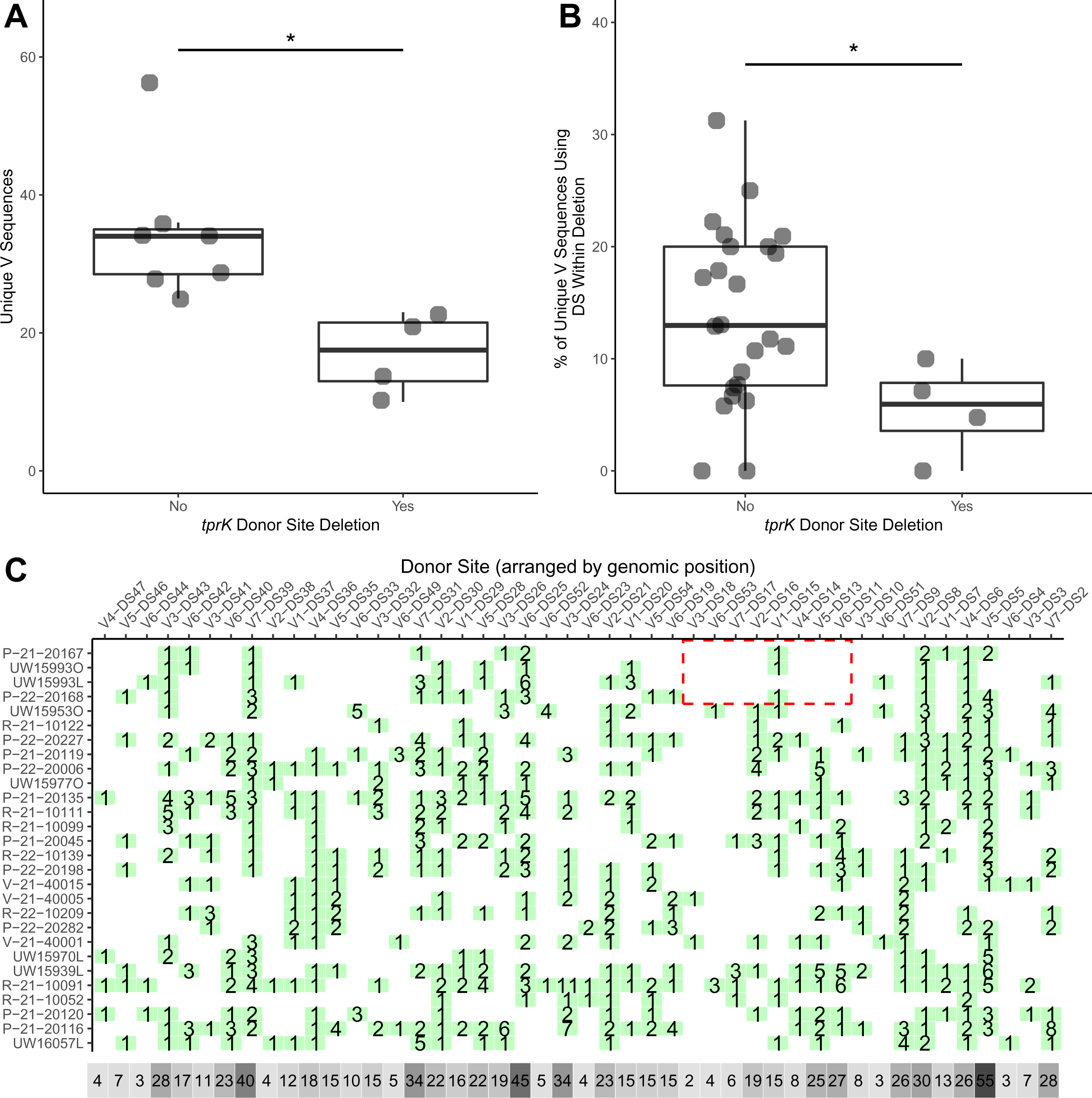
*tprK* variable region composition and donor site usage in TP strains with donor site deletion. A) Among Nichols lineage strains isolated from secondary syphilis lesions, diversity, as measured by the total number of unique V regions detected, is lower in strains bearing a deletion of nine donor sites, *p*<0.05, Welch’s t test. B) As a proportion of total V regions, *tprK* V regions that incorporate the deleted donor sites are less abundant in strains with a deletion of nine donor sites. C) Map of predicted donor site usage by strain. Donor sites are arranged relative to genomic coordinates; x axis is not to scale. Seven donor sites not found in any strain are excluded from the visualization. Green squares indicate the number of *tprK* V region sequences in which each donor site is found. Grey squares indicate the sum of use of each donor site across all samples, with darker grey indicating higher frequency. The deleted region is boxed in red in the affected strains.

We hypothesized that *tprK* V regions recovered from deletion strains would have fewer sequence fragments deriving from the deleted donor sites due to their not being available to generate new variants following adaptive immune-mediated elimination of sequence remnants in place at the time of the deletion. To determine whether deletion of the *tprK* donor sites had an effect on the usage of those donor sites within each *tprK*, we performed a BLAST search using the *tprK* donor site locus as the subject and each *tprK* V region as the query. The top hit for each *tprK* sequence variant was retained, and the pattern of donor site use among samples was calculated. We found that in strains harboring the donor site deletion, the number of unique V regions incorporating the eliminated nine donor sites was significantly lower than in other strains (Figure 5B, *p*<0.05, Welch’s t-test). Indeed, only one of the nine deleted donor sites is present in *tprK* sequences from any of four strains carrying the deletion (Figure 5C).

## Discussion

TP genomic surveillance in the United States has been significantly limited by the availability of molecular testing for diagnosis and relatively low treponemal loads in many patient samples collected for molecular analysis [12]. Here, we offer a preview of a robust end-to-end platform for TP public health genomic surveillance using mostly remnant Aptima swab specimens [26,27]. Our analysis demonstrates the feasibility of using specimens from TMA media for WGS, the correlation between TMA time and quantitative PCR measures of treponemal load, differences in the predominant strains circulating in MSM and MSW/female with syphilis, the ubiquity of azithromycin resistance among circulating TP strains, and the existence of a large deletion in tprK donor site locus associated with reduced tprK diversity.

Our study sheds light on the nature of the syphilis epidemic in Seattle during 2021-2022. In particular, we noted some unique clustering of TP strains by MSM and MSW/women, with one tight MSW/woman SS14 lineage cluster separate from the MSM-associated SS14 lineage and a separate tight MSM-associated Nichols lineage cluster. At the same time, there were a number of unique TP strains in both SS14 and Nichols lineages that had >10 SNVs to phylogenetically closest strain, suggesting greater diversity in the community beyond these two clusters. A large Australian study found an enrichment of MSW and women in a particular SS14 subclade [12]. Previous studies of broader geographic regions have seen some enrichment of MSM individuals in Nichols lineage [15,28], although another study conducted in the Amsterdam area [29] did not confirm this trend. This may be due in part to differences between localities in the populations in which the different lineages circulate, rather than any biological reason. Importantly, whole genome analysis of cultured isolates collected in Seattle between 2001-2011 revealed only two out of 60 (3.3%) belonging to Nichols lineage, with the remainder belonging to the SS14 lineage [13]. Although it is possible that differences in populations sampled during the study periods are driving the shift towards an increased proportion of Nichols-lineage strains, the possibility of a large outbreak of Nichols-lineage syphilis and population expansion among MSM in Seattle is consistent with the very low diversity (median two SNVs) among node B Nichols strains included in this study, and should be explored further. More work and finer sampling is required to understand the degree to which transmission occurs between MSM and MSW/women.

Our data support the use of oropharyngeal or rectal swabs for TP genome recovery, based on the identical genomes recovered. This work extends recently reported TP-MLST data which found identical MLST types among pharyngeal, urogenital, and/or anal swab specimens contemporaneously collected from twelve individuals [31]. Prior work also demonstrated a lack of SNVs within the recoverable portion (54%) of the genome in TP subspecies *pertenue* in banked skin lesion swabs taken two years apart, consistent with the low intra-host diversity of TP genomes seen here [32].

Previous studies have shown no or few *tprK* haplotypes shared among cross-sectional patient samples, gradual accrual of *tprK* V region diversity in immunocompromised hosts and cultured isolates, as well as the requirement for *tprK* donor sites to create this antigenic diversity [7–9, 28]. Here, we showed *tprK* sequence correlates with TP genome relatedness while also serving as a more discriminating locus than the rest of the TP genome. Given the slow evolutionary rate of the TP genome and faster rate of *tprK* gene conversion, it may be worthwhile to include *tprK* deep sequencing data in public health genomic investigations to achieve greater temporal resolution of TP relatedness. Greater *tprK* diversity is associated with secondary syphilis cases [9,30], and *tprK* diversity may be indicative of the time elapsed since infection and helpful for public health investigations. However, correlation with more granular clinical history and epidemiological information is required.

By combining TP genomics with *tprK* profiling, we showed contemporaneous circulation of a TP strain with a significant deletion in the *tprK* donor site locus that was associated diminished *tprK* antigen diversity. Such deletions are relatively rare, as searching of NCBI GenBank recovered one additional MexicoA-like TP strain (MD18Be, NCBI GenBank accession CP073487.2) with a unique 508 bp deletion in the *tprK* donor site locus removing 6 donor sites and creating a novel V7 chimeric donor site [15]. Intriguingly, the *tprK* gene in three of the four isolates in this study, including one sample from all three patients, contained a V1 region comprised entirely of the deleted V1-DS15 donor site. The V1 variable region is enriched in variant sequences comprising a complete donor site sequence and our best interpretation is that this V1 sequence was likely present when the *tprK* donor site locus deletion occurred and only had been recombined out in one of the four specimens [9]. Certainly, more work is required to understand the mechanism of variable region creation both *in vitro* and *in vivo* [10,11,33].

Our study was limited by the lack of known transmission partners to train our TP and *tprK* genomic data. Limited clinical history curtails our current ability to validate *tprK* diversity as a measure of duration of infection. Treponemal genome recovery is limited by the relatively low treponemal loads seen in human specimens, as illustrated by the recovery of complete genomes from fewer than one-third of the TP-positive swabs here, though partial genomes could be recovered from almost one-half of specimens.

We offer a specific molecular testing to TP genome analysis workflow amenable to public health genomic epidemiology for a pathogen with rapidly increasing incidence in the United States and elsewhere. Our data highlight the potential for TP genomics to inform public health epidemiology, as well as the ability of public health genomics to recover variants that bulwark our scientific understanding of this mercurial pathogen.

## Acknowledgements

The authors thank Sheila Lukehart for helpful comments.

## Supplemental Figure Legends

**Supplemental Figure 1. Potential confounders of *tprK* analysis of unique V region sequences recovered.** Variation in *tprK* diversity is not due to A) the number of input genomes (Pearson coefficient 0.204, *p*>0.05, not significant) or B) lineage or clinical stage (Welch’s t test, both *p*>0.05, not significant).

**Supplemental Figure 2. PCR confirmation of *tprK* donor site deletion in deletion-containing and wild-type strains.** PCR and gel electrophoresis was used to confirm the 528 bp deletion detected by whole genome sequencing in strains P-22-20168 and UW15993L. Strains UW15953, P-21-20135, and UW15970L did not have the deletion as detected by whole genome sequencing and were used as controls along with a no template control (NTC).

