## Supplementary figures and images for "Genomic epidemiology of *Treponema pallidum* and circulation of strains with diminished *tprK* antigen variation capability in Seattle, 2021-2022"

### Figure S1

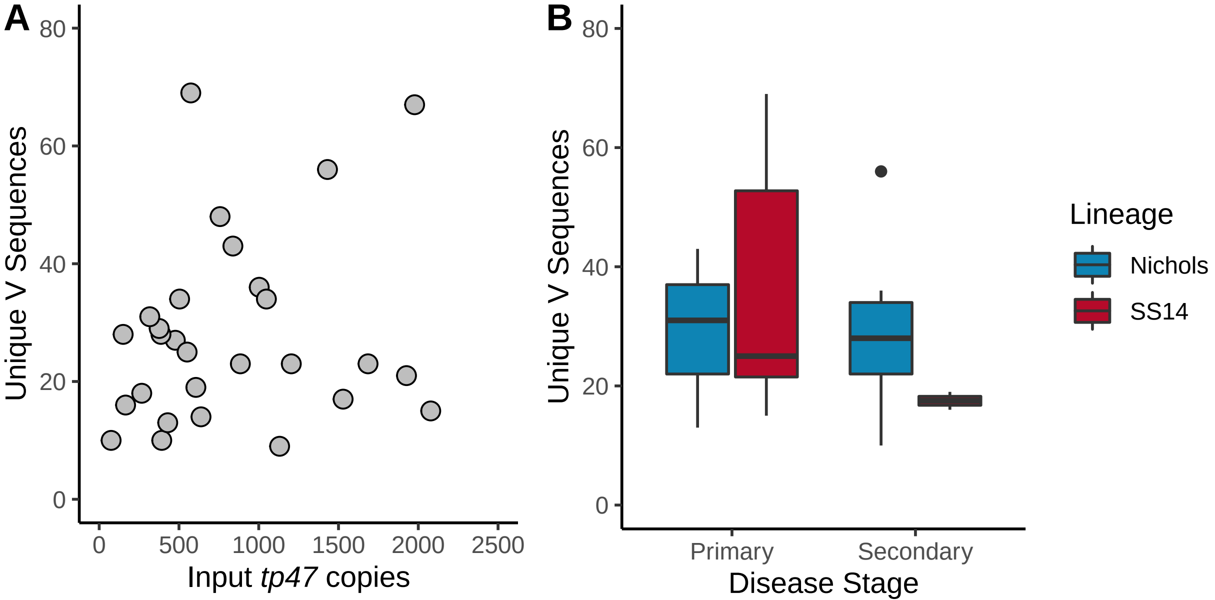

### Figure S2

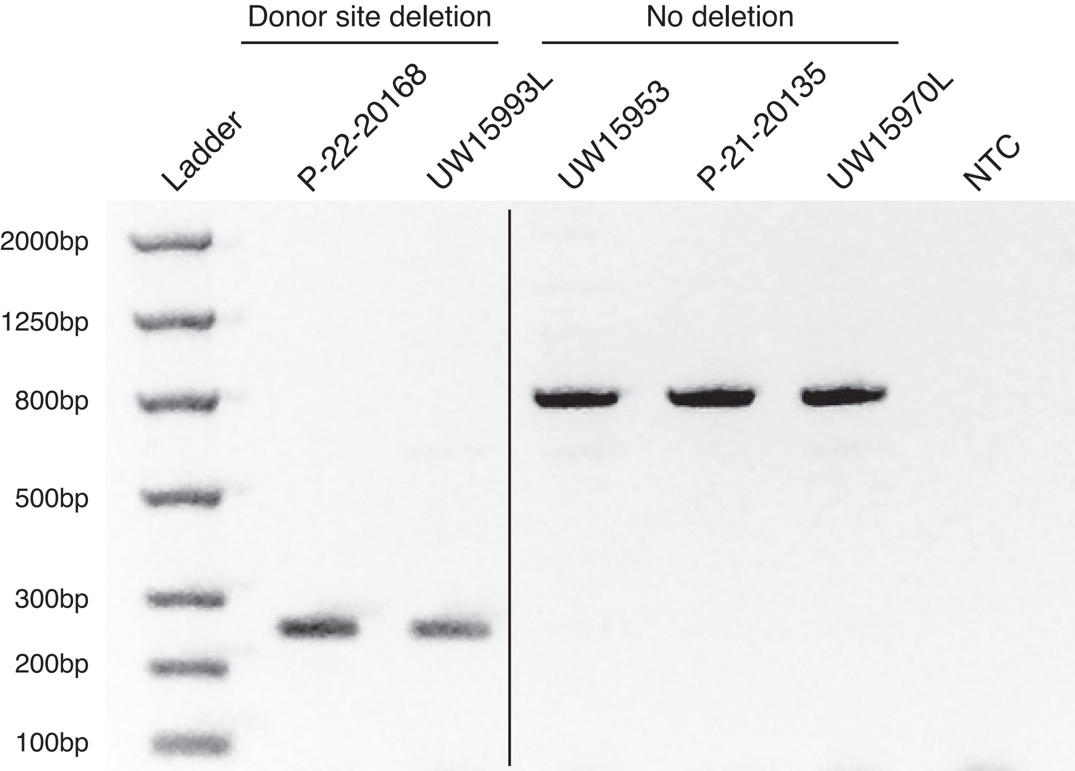
